# DNA methylation at a single cytosine embedded in the W-box *cis*-element repels binding of WRKY transcription factors through steric hindrance

**DOI:** 10.1101/2022.09.26.509487

**Authors:** Magali Charvin, Thierry Halter, Romain Blanc-Mathieu, Pierre Barraud, Magali Aumont-Nicaise, François Parcy, Lionel Navarro

**Affiliations:** Institut de Biologie de l’Ecole Normale Supérieure (IBENS), Centre National de la Recherche Scientifique UMR8197, Institut National de la Santé et de la Recherche Médicale U1024, Paris, France; Laboratoire Physiologie Cellulaire et Végétale, Université Grenoble Alpes, CNRS, CEA, INRAE, IRIG-DBSCI-LPCV, F-38054 Grenoble, France; Expression génétique microbienne, UMR 8261, CNRS, Université Paris Cité, Institut de biologie physico-chimique, IBPC, F-75005 Paris, France; Institute for Integrative Biology of the Cell (I2BC), Université Paris-Saclay, CEA, CNRS, 91198, Gif-sur-Yvette, France

**Author notes:** These authors contributed equally to this work (Magali Charvin, Thierry Halter).

## Abstract

Previously, we showed that the Arabidopsis active demethylase ROS1 prunes DNA methylation at the promoters of a subset of immune-responsive genes to facilitate their transcriptional activation during antibacterial defence (Halter et al., 2021). In particular, ROS1 was shown to demethylate the *RLP43* promoter in a region carrying a functional W-box *cis*-element, thereby ensuring a tight binding of WRKY transcriptional factors (TFs) onto DNA. Here, we first extend these findings by showing that DNA methylation at W-box elements decreases the binding of several Arabidopsis WRKY TFs *in vitro*. Furthermore, we provide evidence that DNA methylation at a single cytosine located in the W-box of the *RLP43* promoter strongly repels DNA binding of an Arabidopsis WRKY TF *in vitro*. Using structural modelling, we demonstrate that this cytosine interacts through van der Waals contacts with the conserved tyrosine of WRKY DNA binding domains. Significantly, our model predicts steric hindrance when a 5-methyl group is present on this specific cytosine, thereby likely preventing tight binding of WRKY DNA binding domains. Finally, because the WRKY motif and the residues involved in DNA contacts are conserved, we propose that this methylation-dependent WRKY-DNA binding inhibitory mechanism must be widespread across plant species.

## INTRODUCTION

Transcription factors (TFs) are central regulators of gene expression and control a wide range of biological processes. They can bind to specific genomic DNA sequences through recognition of Transcription Factor Binding Site (TFBS) and further activate or repress a large repertoire of genes. In eukaryotic cells, transcription is known to be regulated in the context of chromatin, whereby TFs typically compete with nucleosomes for genomic DNA accessibility (Guertin and Lis, 2010; Klemm et al., 2019). Nevertheless, some TFs, referred to as “pioneer” TFs, have the ability to bind nucleosome-rich regions (Clapier and Cairns 2009; Magnani et al., 2011; Iwafuchi-Doi and Zaret, 2014; Iwafuchi-Doi and Zaret, 2016; Zaret et al., 2020; Lai et al., 2021; Jin et al., 2021; Yamaguchi, 2021). Besides nucleosome density, some DNA- or histone-based modifications can additionally modulate the accessibility of TFs on genomic DNA (Klemm et al., 2019). This is notably the case of DNA methylation, a well-characterised epigenetic mark that resides in the methylation of the 5-position of cytosine in DNA, also known as 5-methylcytosine or 5mC. In mammals, almost all 5mC are in the CG context, whereas in plants cytosine methylation occurs in symmetrical (CG or CHG) and in asymmetrical CHH contexts (H = A, T, or C) (Law, J.A & Jacobsen, 2010). In Arabidopsis, methylation nearby a transcriptional start site (TSS) is generally associated with transcriptional gene silencing (Zhang et al., 2006; Ando et al., 2019), suggesting that it restricts DNA/chromatin accessibility for TFs and/or components of the transcription machinery. It has been shown that DNA methylation often blocks TF-DNA binding (Yin et al., 2017; Gaston and Fried, 1995; Watt and Molloy, 1988; Iguchi-Ariga and Schaffner, 1989; Tate and Bird, 1993; Campanero et al., 2000; Comb et al., 1990; O’Malley et al., 2016). For example, a high-throughput TF binding site discovery method, named DNA Affinity Purification sequencing (DAP-seq), reported that ~75% of Arabidopsis TFs (248 out of 327 TFs tested) are sensitive to DNA methylation, meaning that methylation has an inhibitory effect on their DNA binding capacity (O’Malley et al., 2016). A more recent structural and molecular analysis revealed that methylation of the cytosine from the core AAC *cis*-element inhibits DNA-binding of Arabidopsis WEREWOLF (WER) (Wang et al., 2020), a R2R3-MYB TF implicated in root hair patterning (Lee and Schiefelbein, 1999). DNA methylation at the binding site of the Arabidopsis RELATIVE OF EARLY FLOWERING 6 (REF6), a Jumonji C-domain-containing H3K27me3 demethylase, was also shown to repel REF6 binding capacity *in vitro* and *in vivo,* providing further support for a negative impact of methylation on the association of a histone-modifying enzyme onto DNA/chromatin (Qiu et al., 2019). A systematic examination of the effect of methylation on the DNA binding of 542 human TFs also unravelled a large set of TFs that are sensitive to methylation (Yin al., 2017). However, this study also revealed that many human TFs, particularly from the homeodomain family, preferred methylated DNA sites (Yin et al., 2017).

We have recently shown that the Arabidopsis REPRESSOR OF SILENCING 1 (ROS1) actively demethylates the promoter of a subset of defence genes to facilitate their transcriptional activation during antibacterial immunity (Halter et al., 2021). In particular, ROS1 was shown to demethylate the promoter of the Arabidopsis *Receptor Like Protein 43* (*RLP43*) to ensure a proper transcriptional activation of this gene in response to the flagellin-derived peptide flg22 (Halter et al., 2021). Importantly, the *RLP43* promoter DNA region subjected to ROS1-directed demethylation was shown to contain a functional W-box (Halter et al., 2021). This *cis*-regulatory element is the binding motif of WRKYs, a family of plant-specific TFs that play critical roles in the regulation of biotic and abiotic stress signalling (Eulgem et al., 2000; Birkenbihl et al., 2017; Birkenbihl et al., 2018). WRKY TFs all share at least one DNA binding domain of ~60 amino acids, referred to as the WRKY domain, which contains an invariant WRKYGQK sequence that is essential for DNA binding (Maeo et al., 2001). By comparing DAP-seq *versus* ampDAP-seq, in which DNA methylation were removed from the genomic DNA of the Arabidopsis reference accession Columbia-0 (Col-0) by PCR amplification, AtWRKY TFs binding was found globally enriched in the absence of methylation (O’Malley et al., 2016; Halter et al, 2021). A locus-specific DAP-qPCR approach also revealed that the DNA binding of AtWRKY18 and AtWRKY40, two well-characterised flg22-responsive AtWRKYs (Birkenbihl et al., 2017), was detected at the *RLP43* promoter using Col-0 genomic DNA (Halter et al., 2021). By contrast, this binding was almost fully abolished when genomic DNA from *ros1* mutants was used for this assay (Halter et al., 2021). This study therefore demonstrated that the hypermethylation occurring in the *ros1* mutant background at the *RLP43* promoter directly repels the binding of these AtWRKYs *in vitro*. However, the detailed mechanisms responsible for the repelling effect exerted by methylation at the DNA-WRKY interface, and the specific methylcytosine(s) involved in this process, remained elusive.

Here, we used computational, DNA-protein affinity and structural modelling to address this issue. We first show that methylation decreases DNA binding of a subset of Arabidopsis WRKY TFs at their whole targeted genomic regions and at TFBS. Furthermore, we provide evidence indicating that DNA methylation at a single cytosine, located in a W-box element embedded in the *RLP43* promoter, repels WRKY-DNA binding *in vitro*. Finally, we show that the conserved tyrosine of WRKY DNA binding domains interacts with a single cytosine from the W-box, and that the presence of a 5-methyl group at this cytosine alters binding through steric hindrance. Overall, this work describes the first detailed molecular mechanism by which cytosine methylation impedes DNA binding of WRKY TFs.

## RESULTS

### An increased number of methylated cytosines in the whole bound genomic regions and in the TFBS of AtWRKYs reduces their DNA binding affinity

A previous DAP-seq study reported that Arabidopsis WRKY TFs are sensitive to DNA methylation (O’Malley et al., 2016). We also recently demonstrated by DAP-qPCR analysis that the hypermethylation observed in *ros1* mutants at the *RLP43* promoter directly repels DNA binding of WRKY TFs (Halter et al., 2021). This was notably the case of AtWRKY40, a well-characterised flg22-induced WRKY TF that was found to associate both *in vitro* and *in vivo* with the *RLP43* promoter region that is regulated by ROS1 (Halter et al., 2021; Birkenbihl et al., 2017). To get further insights into the mechanisms by which DNA methylation could inhibit AtWRKY40-DNA binding, we made use of available DAP-seq and ampDAP-seq datasets (O’Malley et al., 2016; Lai et al., 2021). We first plotted the DAP/ampDAP signal ratio as a function of the methylation density in the whole AtWRKY40 bound genomic regions, as previously reported for other TFs (Lai et al., 2021). This first analysis revealed that the DAP/ampDAP signal ratio decreased with methylation density (Figure 1A), supporting a negative effect of DNA methylation on AtWRKY40-DNA binding. We further plotted the DAP/ampDAP signal ratio relative to the number of methylated cytosines within WRKY best binding sites, which were identified using position weight matrices in each bound region (Figure 1B; Lai et al., 2021). This analysis revealed that an increased number of methylated cytosines in the TFBS also decreased AtWRKY40-DNA binding. Altered binding to both genomic regions and individual TFBS was also detected for six other AtWRKYs for which DAP/ampDAP datasets were available, namely AtWRKY14, 15, 22, 24, 25 and 27 (Figure 1 – Supplementary Figure 1). Collectively, these data confirm previous findings indicating that DNA methylation inhibits binding of AtWRKYs (O’Malley et al., 2016, Halter et al., 2021). They also show that this binding inhibitory effect is not only detected at the whole bound genomic regions but also at the TFBS recognised by these TFs.

**Figure 1.**
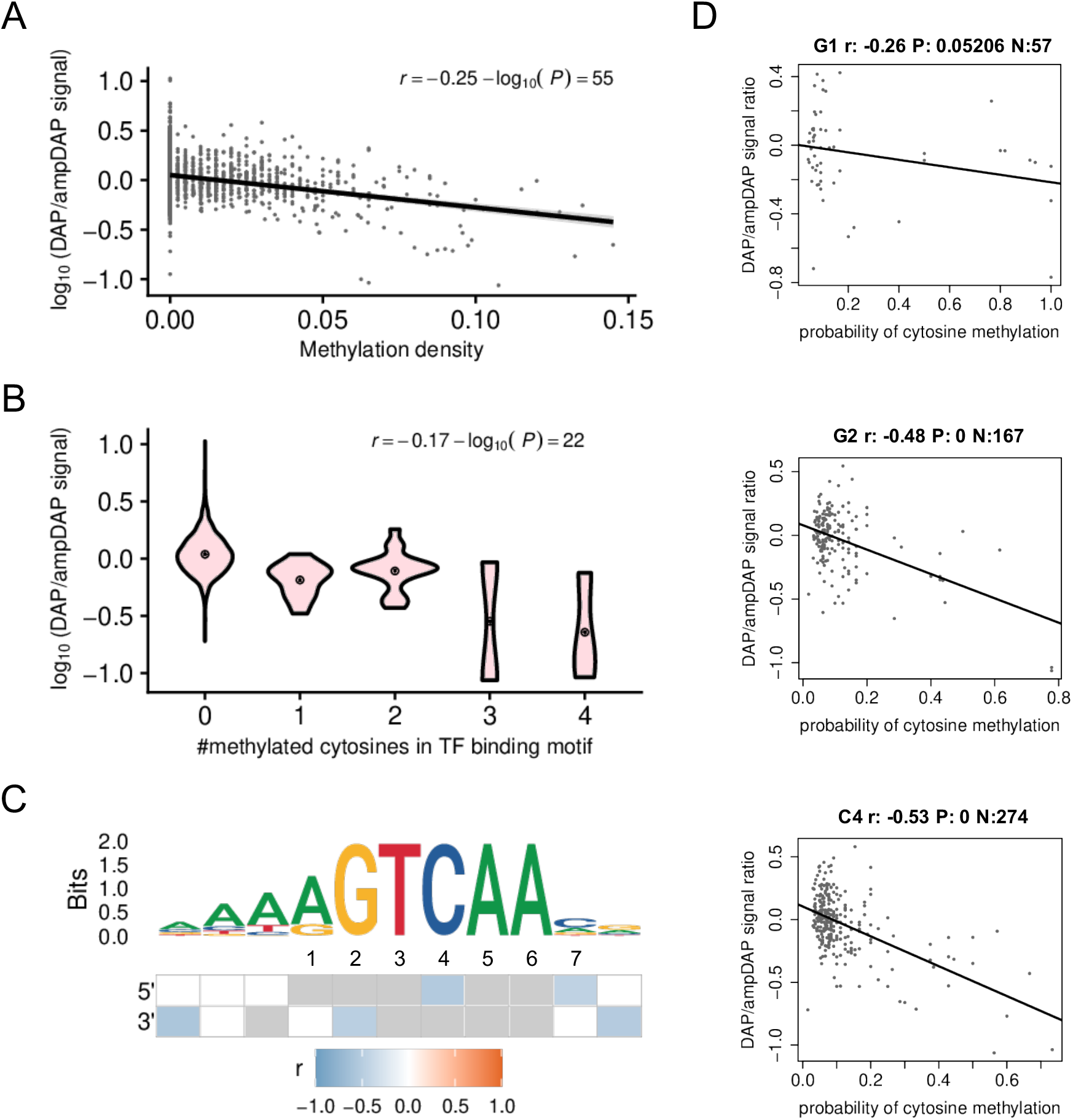
Effect of cytosine methylation on AtWRKY40-DNA binding. (A) Biplot between the DAP/ampDAP signal ratio (peak normalised read coverage in the DAP experiment divided to that in the ampDAP experiment) in a log10 scale and methylation density (proportion of cytosines with a probability of methylation greater than 0.5) within TF bound regions. The increasing methylation density has a negative effect on AtWRKY40 binding. (B) Violin plots of DAP/ampDAP signal ratio in a log10 scale as a function of the number of methylated cytosines in the best TF binding site (TFBS) of each bound region. AtWRKY40 binding is negatively affected by the increased number of methylated cytosines. (C) Binding site sequence motif and the methylation effect on each individual position. The heatmap describes the Pearson’s correlation coefficient (r) between the DAP/ampDAP signal ratio in a log10 scale and the probability of methylation at each position of the best TFBSs. Blank positions have a high false discovery rate (> 5%) and grey indicates positions with less than ten cytosines in the dataset. Correlations are tested on both sides. (D) Effect of methylation on individual positions at the core W-box on AtWRKY40 binding. Relation between methylation probability at a single nucleotide position in the predicted best AtWRKY40 binding site within bound regions, and the log10-scaled relative binding intensity of a DAP-seq *versus* an ampDAP-seq experiment at bound regions for AtWRKY40 at the 3 different cytosine sites. P-values are adjusted for multiple testing using the Benjamini and Hochberg procedure.

### DNA methylation at cytosines embedded in W-box *cis*-elements negatively regulates the binding affinity of AtWRKYs, with 5mC_4_ exhibiting the most significant inhibitory effects

We next used a previously described procedure that computes the impact of individual cytosine methylation on TF-DNA binding by using DAP/ampDAP datasets (Lai et al., 2021). More specifically, we ran this computational approach on the TFBS of the seven Arabidopsis WRKY TFs for which sufficient reads were recovered from the DAP/ampDAP datasets. This analysis revealed that the methylation at different cytosines from the TFBS decreases DNA binding of these AtWRKYs (Figure 1D - Supplementary Figure 2). In particular, DNA methylation at the three cytosines embedded in the W-box motif systematically impacted WRKY-DNA binding, with the most pronounced inhibitory effects detected on cytosines at position 4 on the forward strands (5mC4, CHH context) for six out of the seven AtWRKYs studied (Figure 1D; Figure 1 - Supplementary Figure 2). For example, we found that 5mC_4_, and cytosine methylation at position 2 on the reverse strands (5mC_2_, CHH or CHG contexts), exhibited significant inhibitory effects on AtWRKY40-TFBS binding (r = −0.53 and −0.48, respectively), while methylation at position 1 on the reverse strands (5mC_1_, CHH context) showed milder negative effects (r = −0.26) (Figure 1D). Altogether, these results suggest that methylation at cytosines embedded in the core W-box motif alters AtWRKY-DNA binding, with 5mC_4_ exhibiting the most significant DNA binding inhibitory effects.

### DNA methylation of the cytosine at position 4 of a functional W-box *cis*-element severely reduces AtWRKY40-DNA binding affinity

Although the above computational approach provides useful information on the possible effects of individual cytosine methylation on WRKY-DNA binding, it does not determine the causal role of each cytosine methylation in this process. To address this issue, we used Bio-Layer Interferometry (BLI), a label-free optical technique that measures the interference pattern of white light reflected off a coated biosensor tip (bio-layer) and an internal reference surface. This method is increasingly employed to measure DNA-protein interactions as it can provide binding kinetics in real time, which is not the case of traditional DNA-protein binding assays (e.g. electrophoretic mobility shift assay –EMSA–). In particular, we used BLI to measure the binding capacity of AtWRKY40 at a functional W-box derived from the flg22-responsive *RLP43* promoter (Halter et al., 2021). For this end, we first synthesised two biotinylated single-stranded DNA sequences corresponding to a 16-mer region of the *RLP43* promoter containing the functional W-box, which were either unmethylated or methylated at the cytosine at position 4 on the forward strand of the W-box. The corresponding single-stranded unmethylated or methylated DNA sequences (reverse complementary sequences) were also synthesised and annealed to the above biotinylated sequences to generate DNA duplexes containing either single or combinatorial cytosine methylation at the W-box *cis*-element (Figure 2A). Each DNA duplex was further immobilised onto a streptavidin biosensor, and the resulting bio-layer was introduced into a solution containing different concentrations of a purified truncated AtWRKY40 protein containing its DNA-binding domain (DBD) (Figure 2 – Supplementary Figure 1). We subsequently recorded changes in optical wavelength, which are correlated with the variation in the thickness of the bio-layer resulting from the association of the tested DNA duplex with AtWRKY40 molecules. We found strong wavelength shifts on the bio-layer containing the unmethylated DNA duplex, a feature that was observed at the five AtWRKY40 protein concentrations tested (Figure 2B). These results demonstrate effective interactions between AtWRKY40 proteins and the unmethylated DNA duplex, which were further supported by a constant of dissociation (K_D_) value of 630 nM at the steady state level (Figure 2C). We next analysed the effect of DNA methylation at each cytosine of the W-box on the ability of AtWRKY40 to bind the DNA duplex. We found that methylation at the two cytosines, namely 5mC_1_ and 5mC_2_ (both in CHH contexts), located on the reverse strand of the W-box did not alter DNA binding, as wavelength shifts and K_D_ value were similar to the ones detected with the unmethylated DNA duplex (Figure 2). By contrast, the wavelength shifts were significantly reduced, and the K_D_ value significantly higher (12 μM), when the cytosine at position 4 from the forward stand (5mC_4_, CHH context) of the W-box was methylated (Figure 2). When the same analysis was conducted side by side with a cytosine to thymine substitution at C4, which is known to abolish WRKY-DNA bindings (Ciolkowski et al., 2008), we found that the binding inhibitory effect triggered by 5mC_4_ was almost as strong as the one caused by this point mutation (Figure 2 – Supplementary Figure 2). This result further supports a severe, albeit not complete, negative impact of 5mC_4_ on AtWRKY40-DNA binding. Furthermore, no additive effects on the wavelength shift, nor on the K_D_ value (9.2 μM), were found when the C1 and C2 nucleotides from the reverse strand of the W-box were methylated besides 5mC_4_ (Figure 2). Altogether, these *in vitro* data provide solid evidence that 5mC_4_ at a functional W-box *cis*-element has a strong and specific inhibitory effect on the ability of AtWRKY40 to bind DNA.

**Figure 2.**
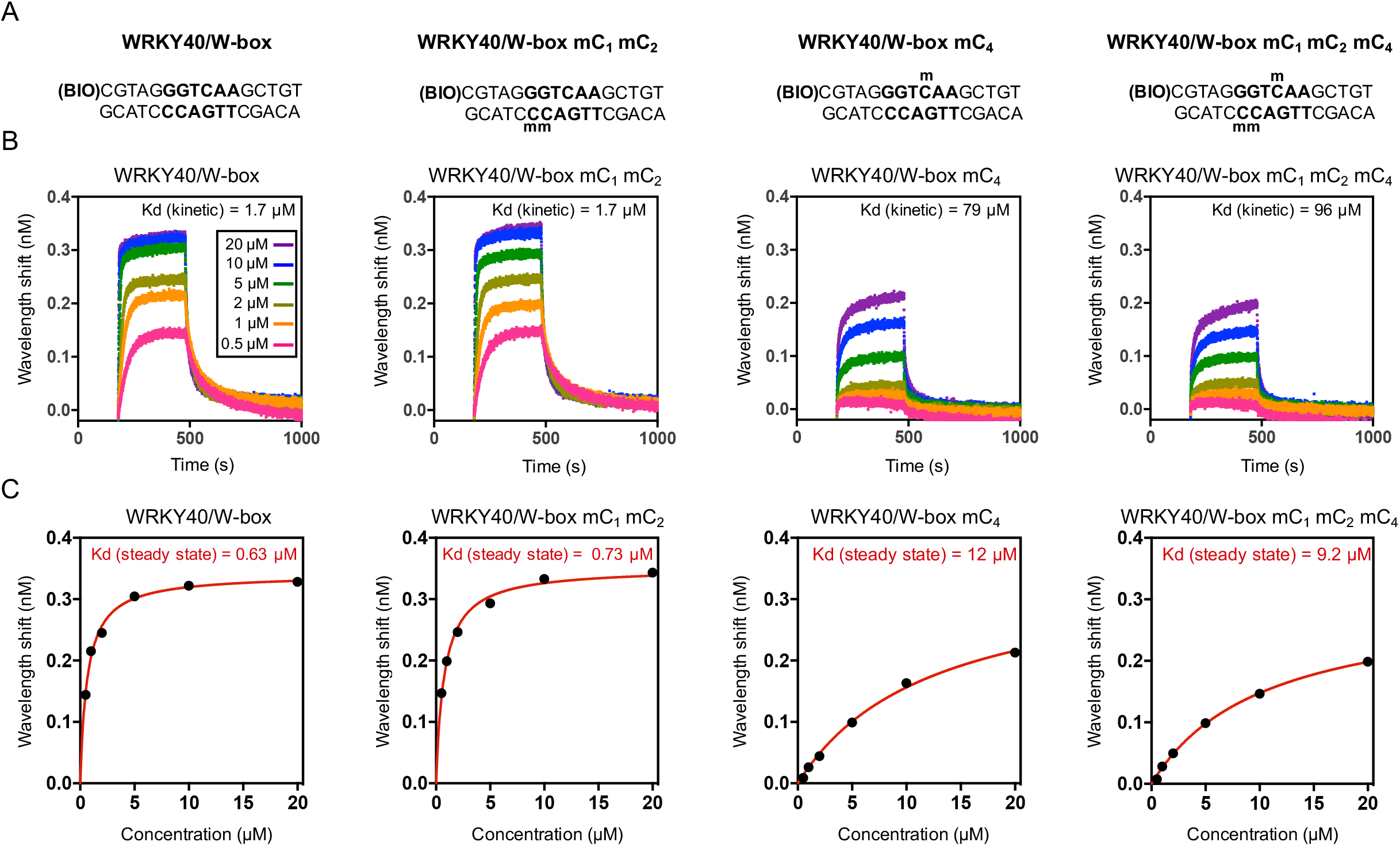
Methylation at the cytosine C4 embedded in a functional W-box has a strong negative impact on AtWRKY40-DNA binding. (A) Biotinylated DNA duplexes used for the BLI experiments. (B) Binding curves of AtWRKY40 DBD with DNA duplexes containing W- boxes with indicated methylation status. (C) BLI-derived steady state analysis representing binding responses of AtWRKY40 DBD (nM) to DNA duplexes as a function of AtWRKY40 DBD concentration.

### The tyrosine residue of the WRKY domains of AtWRKY40 and AtWRKY4 make van der Waals contacts with the cytosine at position 4 of the core W-box *cis*-element

To further understand the detailed mechanism by which 5mC_4_ from the W-box repels AtWRKY40-DNA binding, we sought to build a structural model of the AtWRKY40 DBD in complex with a W-box DNA duplex. For this end, we first generated a structural model of the WRKY domain of AtWRKY40 (residues 131-207) using homology modelling with Swiss-Model (Waterhouse et al., 2018; Figure 3 – Supplementary Figure 1 A, B). We then built the protein-DNA complex model, by superimposing the AtWRKY40 homology model onto the Nuclear Magnetic Resonance (NMR) structural ensemble of the C-terminal WRKY domain of AtWYRK4, in complex with a W-box DNA element (PDB code 2lex) (Yamasaki et al., 2012). The resulting bundle of structures was refined with a restrained simulated annealing protocol as described in material and methods. The 10 lowest energy structures of the protein-DNA complex were pooled (Figure 3 – Supplementary Figure 1D) and protein-DNA contacts were carefully analysed in this bundle (Figure 3). Importantly, we noticed that in all structures of the bundle, the aromatic ring of C4 makes van der Waals contacts with the side-chain of Y154 (Figure 3B, D). Because our model is based on the structure of AtWRKY4 in complex with a core W-box element (Yamasaki et al., 2012), we also analysed the interface between AtWRKY4 and the W-box DNA. We found that the binding mode of AtWRKY4 also involves van der Waals contacts between the aromatic ring of C4 in the W-box and the Y417 residue, the equivalent of Y154 in AtWRKY40 (Figure 3 – Supplementary Figure 2). These analyses therefore unveiled the presence of van der Waals interactions between the tyrosine residues of the WRKY domains of two Arabidopsis WRKYs and the C4 of the W-box DNA.

**Figure 3.**
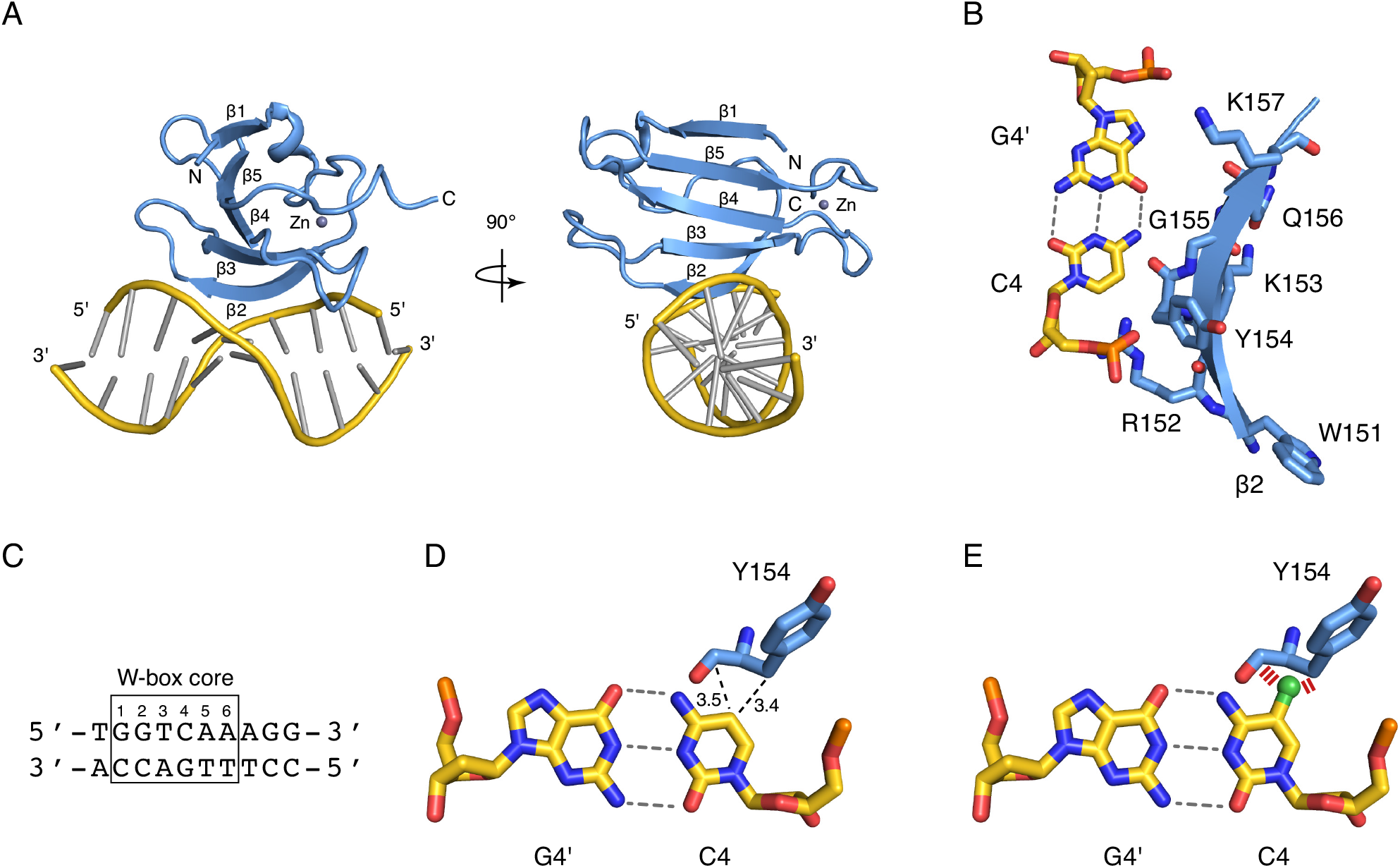
Structural model of the AtWRKY40 DBD in complex with a W-box DNA duplex. (A) Lowest energy structure from the ensemble of models of the complex between AtWRKY40 and the W-box element of panel C. The protein is shown as a cartoon in light blue, the five β-strands are labelled, and the zinc ion is shown as a grey dot. The DNA backbone is shown as a cartoon in yellow and the base-pairs are shown with the ladder representation in grey. (B) Close-up view of the WRKY motif of β2 (W151-K157). The protein is shown in light blue with the side-chains represented as sticks. The C4-G4’ DNA base-pair is shown as sticks in yellow. The β2 strand enters deeply into the DNA major groove at the level of the C4-G4’ base-pair. In particular, the aromatic ring of C4 makes van der Waals contacts with the side-chain of Y154. (C) Sequence of the DNA W-box element used for modelling the interaction of AtWRKY40 with DNA. (D) The position 5 of C4 makes van der Waals contacts with Y154. (E) Modelling a methyl group (in green) onto unmodified C4 from the W-box core reveals steric hindrance (indicated with red strips) with Y154.

### The interaction between the tyrosine residues of the WRKY domains of AtWRKY40 and 4 and the C4 of the W-box is incompatible with a 5-methyl group at this nucleotide

We next modeled the impact of 5mC_4_ on the binding of the C4 nucleotide to the Y154 and Y417 residues of AtWRKY40 and AtWRKY4, respectively. Importantly, our model predicts that the presence of a 5-methyl group at this specific cytosine would prevent the β2-strand of the DNA-binding domains of both AtWRKY TFs to deeply enter into the DNA major groove at the level of the W-box element (Figure 3A; Figure 3 – Supplementary Figure 2). This interaction mode is therefore incompatible with the presence of a 5-methyl group on this cytosine (Figure 3E; Figure 3 – Supplementary Figure 2). In addition, we found that both the WRKY motif in β2, and the residues involved in DNA contacts, are well conserved among different WRKY domains (Figure 3 – Supplementary Figure 3). The steric hindrance preventing optimal W-box DNA-binding by AtWRKY40 and 4 in the presence of a 5-methyl group (Figure 3E; Figure 3 – Supplementary Figure 2), is thus likely a feature shared by several WRKY domains. Overall, our model suggests that 5mC_4_ of the core W-box *cis*-element repels binding of WRKY TFs by preventing the occurrence of van der Waals interactions between the C4 nucleotide of the W-box and the conserved tyrosine residue of WRKY DNA binding domains.

## DISCUSSION

In all organisms, the regulation of TF-DNA binding plays a central role in fine-tuning gene expression, and in turn governing biological functions. One layer of such regulation is directed by DNA methylation, an epigenetic mark that can negatively or positively control TF-DNA binding (Yin et al., 2017; Gaston and Fried, 1995; Watt and Molloy, 1988; Iguchi-Ariga and Schaffner, 1989; Tate and Bird, 1993; Campanero et al., 2000; Comb et al., 1990; O’Malley et al., 2016). In humans, early studies conducted on individual loci demonstrated inhibitory effects of DNA methylation on the binding of several TFs (Gaston and Fried, 1995; Watt and Molloy, 1988; Iguchi-Ariga and Schaffner, 1989; Tate and Bird, 1993; Campanero et al., 2000; Comb et al., 1990). Using methylation sensitive SELEX (Systematic Evolution of Ligands by Exponential Enrichment), a more recent study broadened these findings by showing that a large set of human TFs are sensitive to DNA methylation (Yin et al., 2017). These TFs mostly belong to bHLH, bZIP and ETS families, and the repelling effects exerted by cytosine methylation at their target sequences were found to be caused by steric hindrance (Yin et al., 2017). Intriguingly, many other human TFs, particularly from the homeodomain family, preferred methylated cytosines (Yin et al., 2017). These TFs were found to exhibit a preference for methylated cytosines through direct hydrophobic interactions with the 5-methyl group of methylcytosines (Yin et al., 2017). In Arabidopsis, a systemic examination of the effect of methylation on TF-DNA binding has also been conducted (O’Malley et al., 2016). This DAP-seq analysis revealed that ~75% of the Arabidopsis TFs studied were sensitive to DNA methylation, while only ~5% preferred methylation (O’Malley et al., 2016). This study therefore indicates that the DNA binding of Arabidopsis TFs is mostly negatively regulated by methylation through poorly defined mechanisms.

WRKY proteins form one of the largest plant TF families that has been implicated in various processes, including growth, development and stress signalling (Hinderhofer and Zentgraf, 2001; Johnson et al., 2002; Luo et al., 2005). The most extensively characterised function of WRKY TFs resides in the regulation of abiotic and biotic stress signalling (Ülker and Somssich, 2004; Eulgem and Somssich, 2007). Seminal studies on the mode of action of WRKYs have demonstrated that these TFs bind to the core W-box *cis*-element (TTGAC/T or G/ATCAA), which represents the minimal consensus motif required for specific DNA binding (de Pater et al., 1996; Ruhston et al., 1996; Wang et al., 1998; Chen and Chen et al., 2000; Cormack et al., 2002; Ulker and Sommssich 2004; Eulgem and Somssich 2007). Other studies provided evidence that each amino acid residue from the WRKY domain plays a central role in WRKY-DNA binding (Maeo et al., 2001; Duan et al., 2007; Ciolkowski et al., 2008). Accordingly, non-synonymous mutations in any of the amino acid from the WRKY domain, or substitutions in each of the nucleotide forming the core W-box sequence, fully abolish the ability of WRKY TFs to bind DNA (Ruhston et al., 1996; Chen and Chen et al., 2000; Cormack et al., 2002; Ulker and Sommssich 2004; Eulgem and Somssich 2007; Maeo et al., 2001; Duan et al., 2007; Ciolkowski et al., 2008). Here, we have conducted an in-depth characterization of the impact that cytosine methylation could have on WRKY-DNA binding. Using DAP/ampDAP-seq datasets, we first showed that the DNA methylation density inhibited the binding of seven Arabidopsis WRKYs on their targeted genomic regions (Figure 1). This supports previous findings demonstrating that methylation inhibits WRKY-DNA binding (O’Malley et al., 2016, Halter et al., 2021). Furthermore, we showed that an increased number of methylated cytosines at TFBS negatively regulates the binding of these AtWRKYs (Figure 1). Using a recently described computational approach on DAP/ampDAP-seq datasets (Lai et al., 2021), we further identified individual methylated cytosines from the WRKY TFBS that inhibit DNA binding (Figure 1). In particular, we found that the methylation of cytosines located in the core W-box *cis*-element had repelling effects on the DNA binding of the different AtWRKYs (Figure 1C/D, Figure 1 – Supplementary Figure 1). Importantly, we found that 5mC_4_ exhibited the most pronounced inhibitory effect on the binding of the different AtWRKY TFs studied, with the exception of AtWRKY22 (Figure 1C/D, Figure 1 - Supplementary Figure 1 and 2). Furthermore, we noticed that some methylated cytosines surrounding the core W-box elements from the TFBS also exhibited inhibitory effects on WRKY-DNA binding (Figure 1; Figure 1 - Supplementary Figure 1). This was for instance the case of 5mC_7_, which systematically showed reduced binding of different AtWRKY TFs (Figure 1; Figure 1 - Supplementary Figure 1). Collectively, these observations suggest that the methylation environment into which the W-box is embedded might additionally influence WRKY-DNA binding. Because the core W-box sequences are conserved across WRKY target sequences (Eulgem et al., 2000), the number and methylation status of the surrounding cytosines might represent relevant features driving the binding selectivity of WRKY TFs. This would add another epigenetic-based layer of regulation compared to the classical nucleotide composition of the sequences neighbouring the W-box, which has already been proposed to contribute to binding specificity (Ciolkowski et al., 2008).

In both plants and mammals, active DNA demethylation fine-tunes the expression of genes involved in developmental processes and stress signalling (Deleris et al., 2016; Yu et al., 2013; Dowen et al., 2012; Halter et al., 2021; Khouider et al., 2021; Yamamuro et al., 2014; Kim et al., 2019). In particular, there is emerging evidence indicating that active DNA demethylation controls the expression of immune-responsive genes in the context of host-pathogen interactions (Deleris et al., 2016; Yu et al., 2013; Dowen et al., 2012; Halter et al., 2021; Pacis et al., 2015; Pacis et al., 2019; Lopez Sanchez et al., 2016; Le et al., 2014; Schumann et al., 2019; Huang et al., 2022). In human cells, earlier studies suggested a role for active demethylation in modulating the DNA/chromatin accessibility of TFs during bacterial infection or elicitation (Pacis et al., 2015). For example, a large set of genomic regions were found demethylated in human dendritic cells infected with *Mycobacterium tuberculosis*, leading to the chromatin relaxation at enhancers carrying stress-responsive *cis*-regulatory elements (Pacis et al., 2015). A more recent follow-up study suggested that active demethylation is unlikely required for the DNA/chromatin-based recruitment of TFs, but is more likely occurring as a consequence of TFs binding to these genomic regions (Pacis et al., 2019). However, the exact role that demethylation could play in the control of TF-DNA binding remains ambiguous, because this work has not been conducted in human cells lacking active DNA demethylases. In Arabidopsis, a seminal work showed that the active demethylase ROS1 facilitates the flg22-induced transcriptional activation of the nucleotide binding leucine-rich repeat immune receptor *RESISTANCE METHYLATED GENE 1* (*RMGT1*), presumably by erasing cytosine methylation in a promoter region carrying W-box *cis*-regulatory elements (Yu et al., 2013). More recently, a causal effect of ROS1-directed demethylation on the binding of AtWRKYs to a discrete promoter region of the surface immune receptor *RLP43* has been demonstrated (Halter et al., 2021). More specifically, the hypermethylation detected in the *ros1* mutant background at the *RLP43* promoter was found to repel the binding of AtWRKY18 and AtWRKY40, two well-characterised flg22- responsive WRKY TFs (Halter et al., 2021; Birkenbihl et al., 2017). However, the detailed mechanism by which the hypermethylation at this *RLP43* promoter region impinges WRKY-DNA binding in the absence of ROS1 remained elusive. Here, we assessed whether cytosine methylation in the ROS1-targeted *RLP43* promoter region carrying a functional W-box *cis*-element could interfere with AtWRKY40-DNA binding. Using a BLI approach, we demonstrated that the methylation at the cytosine at position 4 (5mC_4_) of this W-box severely reduced the ability of AtWRKY40 to bind DNA, an effect that was almost as strong as the one triggered by a point mutation in this specific cytosine (Figure 2, Figure 2 – Supplementary Figure 2). By contrast, methylation of the other cytosines, located on the opposite strand of this W-box sequence, did not trigger any repelling effect on AtWRKY40-DNA binding (Figure 2). Collectively, these data support a strong and specific repelling effect of 5mC_4_ on the binding of AtWRKY40 to this W-box DNA motif. They also further confirm, through an independent and complementary experimental approach to DAP/ampDAP analysis, that 5mC_4_ exerts a potent inhibitory effect on AtWRKY40-DNA binding. However, it is noteworthy that 5mC_4_ did not fully abolished TF binding, as previously observed when the whole promoter-regulatory region of *RLP43* was hypermethylated (Halter et al., 2021). This suggests that the methylation of cytosines flanking this functional W-box must additionally contribute to the DNA binding inhibitory effect (as discussed in the previous section). Using structural modelling, we further showed that the cytosine at position 4 of the core W-box element makes van der Waals contacts with the tyrosine residue of the WRKY domain of AtWRKY40 (Figure 3). Our model also predicts that the negative effect triggered by 5mC_4_ at the W-box is caused by steric hindrance, that likely prevents the β2-strand of AtWRKY40 WRKY domain to deeply enter into the DNA major groove at the level of the W-box element, thereby preventing tight binding to DNA (Figure 3). This prediction was also true when we analysed the interface between AtWRKY4, whose structure was used to build our AtWRKY40 model, and the W-box DNA (Figure 3 – Supplementary Figure 2). These observations suggest that steric hindrance caused by cytosine methylation is a general phenomenon that does not only dampen DNA binding of human TFs, as previously shown (Yin et al., 2017), but also of plant TFs (Wang et al., 2020; this study). In addition, we found that the WRKY motifs of the above AtWRKYs in β2, and residues involved in DNA contacts, were well conserved among different WRKY domains (Figure 3 – Supplementary Figure 3). Based on these data, and on previous studies reporting a strong conservation of the WRKY domains and of the core W-box motifs (Eulgem et al., 2000), we propose that this methylation-dependent binding inhibitory mechanism must operate in all plant species expressing WRKY TFs.

## MATERIAL AND METHODS

### Genome wide effects of methylated cytosines on WRKY-DNA binding

To assess the effect of cytosine methylation on the WRKY binding, we compared the binding intensity of AtWRKYs in a DNA Affinity Purification sequencing (DAP-seq) experiment and in an amplified DAP-seq experiment. In the latter experiment, methylation marks are erased during PCR-based amplification. We tested the correlation between the DAP/ampDAP signal ratio and the methylation levels at all bound regions, at the best TF binding site (TFBS) in the bound region, and at each position of the TFBS as previously described (Lai et al, 2021). In brief, DAP-seq and ampDAP-seq datasets of WRKY TFs (O’Malley et al, 2016) were mapped against the Arabidopsis TAIR10 genome assembly using bowtie2 (Langmead and Salzberg, 2012). Resulting alignment files were used to identify genomic regions bound by the factor (so called peaks) using MACS2 (https://www.biorxiv.org/content/10.1101/496521v1) and MSPC (Jalili et al., 2018). Binding intensity at each bound region is expressed as number of reads mapped per million of reads mapped in total bound regions. TFBSs were searched in bound regions using a position weight matrix (PWM) constructed for each TFs using MEME (Machanick and Bailey, 2011). The probability of cytosine methylation was taken from (Zhang et al., 2016). Methylation density (the number of methylated cytosines in a bound region) was defined as the number of cytosines with a probability of methylation greater than 50%. Association between the relative binding intensity and methylation levels was assessed using Pearson’s correlation test from R package “stats”.

### Production of recombinant AtWRKY40 DBD

DBD AtWRKY40 was cloned in the pET28a destination vector that carries a 6His-tag in N-terminal following classical digestion/ligation with NdeI and XhoI enzymes and T4 DNA ligase and expressed in the *E. coli* strain BL21(DE3) codon plus (ThermoFisher, EC0114) in 1L of Terrific broth (TB) (Supplementary file 1). Cultures were grown at 37°C until they reached an OD600 of 2 and protein production was induced with 1 mM Isopropyl-β-D-thiogalactopyranoside (IPTG), followed by overnight growth at 18°C. Bacterial cells expressing WRKY domain of AtWRKY40 were collected by centrifugation, resuspended in Lysis Buffer (1.5X PBS, 1 mM MgAc_2_, 0,1 % NP-40, 20mM imidazole, 10 % glycerol), and lysed by sonication during 4 min on ice. Lysate was clarified by high-speed centrifugation (18000 rpm) and then purified on 250 μL of Ni-NTA resin (Thermo Fisher Scientific, 88221). Resin was pre-equilibrated in Lysis Buffer, and supernatant was incubated with resin for 2h at 4°C. His fusion proteins linked to the resin were washed once with Lysis Buffer, then once with Wash Buffer (1.5X PBS, 250 mM NaCl, 1 mM MgAc_2_, 0,1 % NP-40, 50 mM imidazole, 10 % glycerol), and finally with Lysis Buffer. Proteins were eluted in Elution Buffer (1.5X PBS, 1 mM MgAc2, 0,1 % NP-40, 150 mM imidazole, 10 % glycerol). Excess imidazole was removed by overnight dialysis using Spectrum™ Labs Spectra/Por™ 2 12-14 kD MWCO (FisherScientific, 15310762) into Dialysis Buffer (1.5X PBS, 1 mM MgAc_2_, 10 % glycerol, 2 mM DTT) before storage at −80°C. Protein extracts recovered at different steps of the purification procedure were resolved by SDS-PAGE on a 15% acrylamide gel, which was stained with Coomassie blue. A band above 15 kDa, corresponding to AtWRKY40 DBD, was clearly visible (Figure 2 – Supplementary Figure 1).

### Measurement of AtWRKY4O-DNA interaction by Bio-Layer Interferometry

BLI experiments were conducted using a FortéBio’s Octet® RED96e system (Sartorius) and Streptavidin (SA) Biosensors. BLI monitors wavelength shifts (nm) resulting from changes in the optical thickness of the sensor surface during association or dissociation of the analyte. All BLI experiments were performed at 25 °C under 1000 rpm stirring. The streptavidin biosensor was hydrated in a 96-well plate containing buffer (50mM Hepes pH7, 150mM NaCl) for at least 10 min before each experiment. The biotinylated oligonucleotide (50μM) was annealed to its non-biotinylated reverse complementary oligonucleotide (60μM) in an Hepes-NaCl buffer (Supplementary file 1). The mix was heated up to 95°C and hybridization occurred during overnight cooling to 25°C. DNA duplexes at 40 nM were immobilised in buffer onto the surface of the SA biosensor through a cycle of Baseline (120 s), Loading (300 s), and Baseline (120 s). The sensors were immobilised one by one in order to obtain the same immobilisation rate (0.25 nm). A reference SA biosensor was prepared in parallel without biotinylated primer immobilised. The DBD of AtWRKY40 was diluted to the corresponding concentrations (10 μM, 5 μM, 2 μM, 1 μM, 0.5 μM, 0.25 μM) in running buffer (50 mM Hepes pH7, 150 mM NaCl, 0.05% Tween20). Protein-DNA interactions were then monitored during 300 s in wells containing 200 μL samples of AtWRKY40 proteins at each concentration. At the end of each binding step, the sensors were transferred into a protein-free binding buffer to follow the dissociation kinetics for 900 s. Data were analysed using FortéBio Data Analysis 12.2 (Sartorius, FortéBio®). The resulting data were fitted into a 1:1 binding model from which K_on_ and K_off_ values were obtained, and then the equilibrium dissociation constant K_D_ values were calculated.

### Modelling the interaction with the W-box DNA

A homology model of the WRKY domain of AtWRKY40 was generated using Swiss-Model (Waterhouse et al., 2018), using the C-terminal WRKY domain of AtWRKY1 as a template (PDB code 2ayd) (Duan et al., 2007). The homology model of AtWRKY40 was then superimposed on the backbone atoms with each of the 20 refined structures of AtWRKY4-C in complex with a W-box DNA element (PDB code 2lex) (Yamasaki et al., 2012). This initial superposition allowed us to generate 400 starting structures of protein/DNA complexes between AtWRKY40 and the W-box DNA, each of them built as a unique pair of conformers. Each individual structure was subjected to a refinement protocol with no experimental energy terms in CNS 1.21 (Brunger et al., 2007), following previously described procedures (Barraud et al., 2014; Barraud et al., 2012). First, the structures were energy minimised with a conjugate gradient minimization, and subsequently a rigid body minimization with two rigid groups defined as one for the protein and one for the DNA. Second, these minimised structures were subjected to a restrained simulated annealing protocol in implicit water. It consisted of 6 ps of dynamics at 1000 K followed by cooling to 25 K over 26 ps. Different types of restraint were applied for the interface and for the rest of the molecules; (i) the side chains of beta-strands 2 and 3 (152-157 and 166-170) were set to unrestrained atoms; (ii) the backbone of beta-strands 2 and 3 (152-157 and 166-170) were harmonically restrained to their initial position, allowing small motions for these parts; (iii) all the rest of the protein and the DNA were set to fixed atoms. The resulting complexes were finally energy minimised and the 10 best energy structures were pooled as the refined ensemble and analysed with PYMOL.

## Acknowledgements

This work was supported by the French funding research agency “L’Agence nationale de la recherche” ANR-18-CE20-0020-NEPHRON (to L.N.), ANR-17-CE20-0014-01-UBIFLOR (to F.P.). This work has also benefited from the PIM Facility of I2BC supported by French Infrastructure for Integrated Structural Biology (FRISBI) ANR-10-INBS-05. We also thank members of the Navarro Lab for the critical reading of the manuscript.

## Author contributions

T.H., P.B., L.N. designed research; M.C., R.B-M, P.B., M A-N performed research, R.B-M., F.P. contributed to new bioinformatic analytic tool; M.C, T.H., R.B-M, P.B., L.N. analysed data; and T.H., L.N. wrote the paper.

---

**Figure 1 - Supplementary Figure 1.**
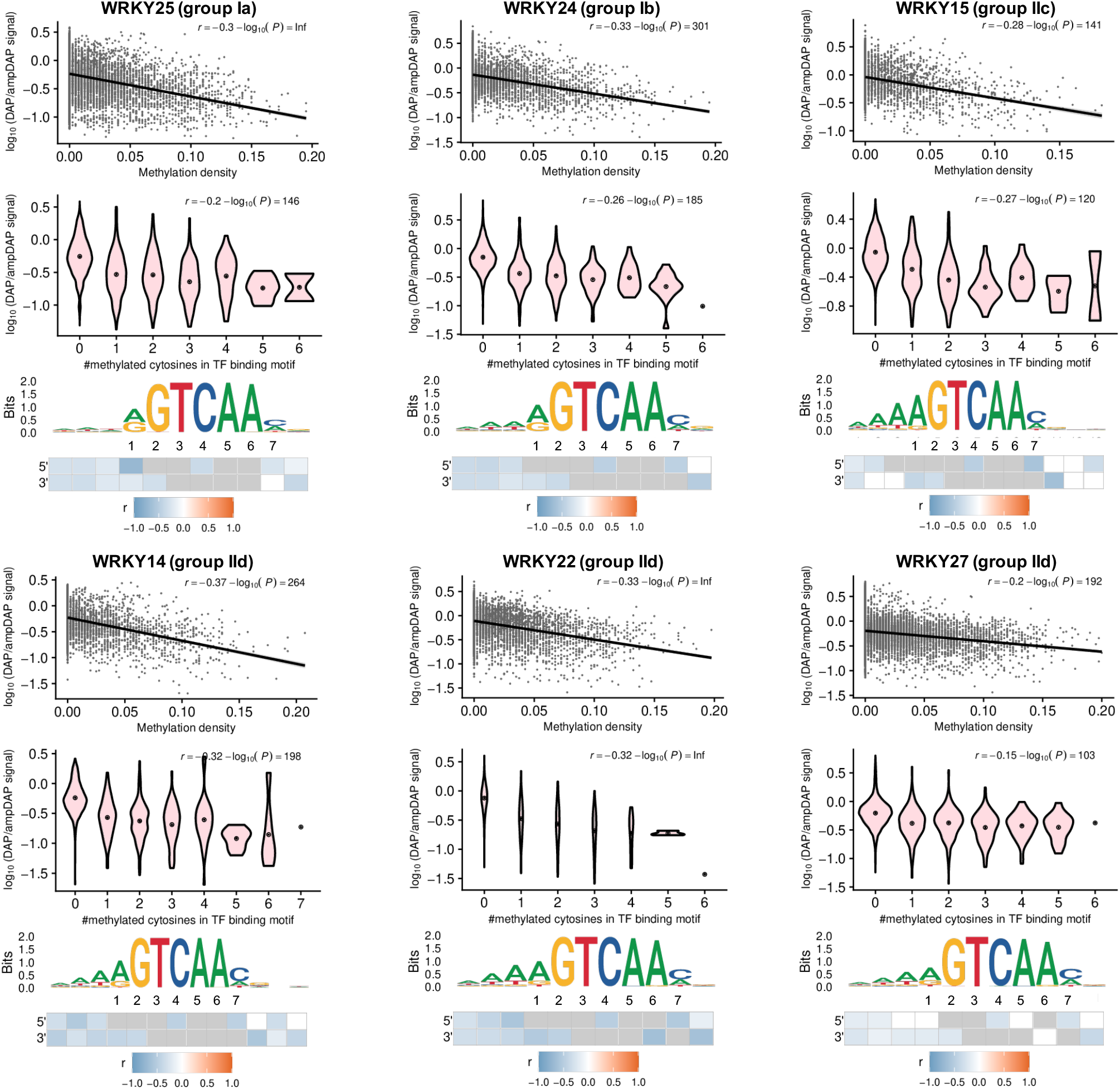
Effect of methylation on the DNA binding of At-WRKYs. The impact of methylation density (upper panel), of the number of methylated cytosines (middle panel) on WRKY binding and individual cytosines (lower panel) was tested for several AtWRKY TFs.

**Figure 1 - Supplementary Figure 2.**
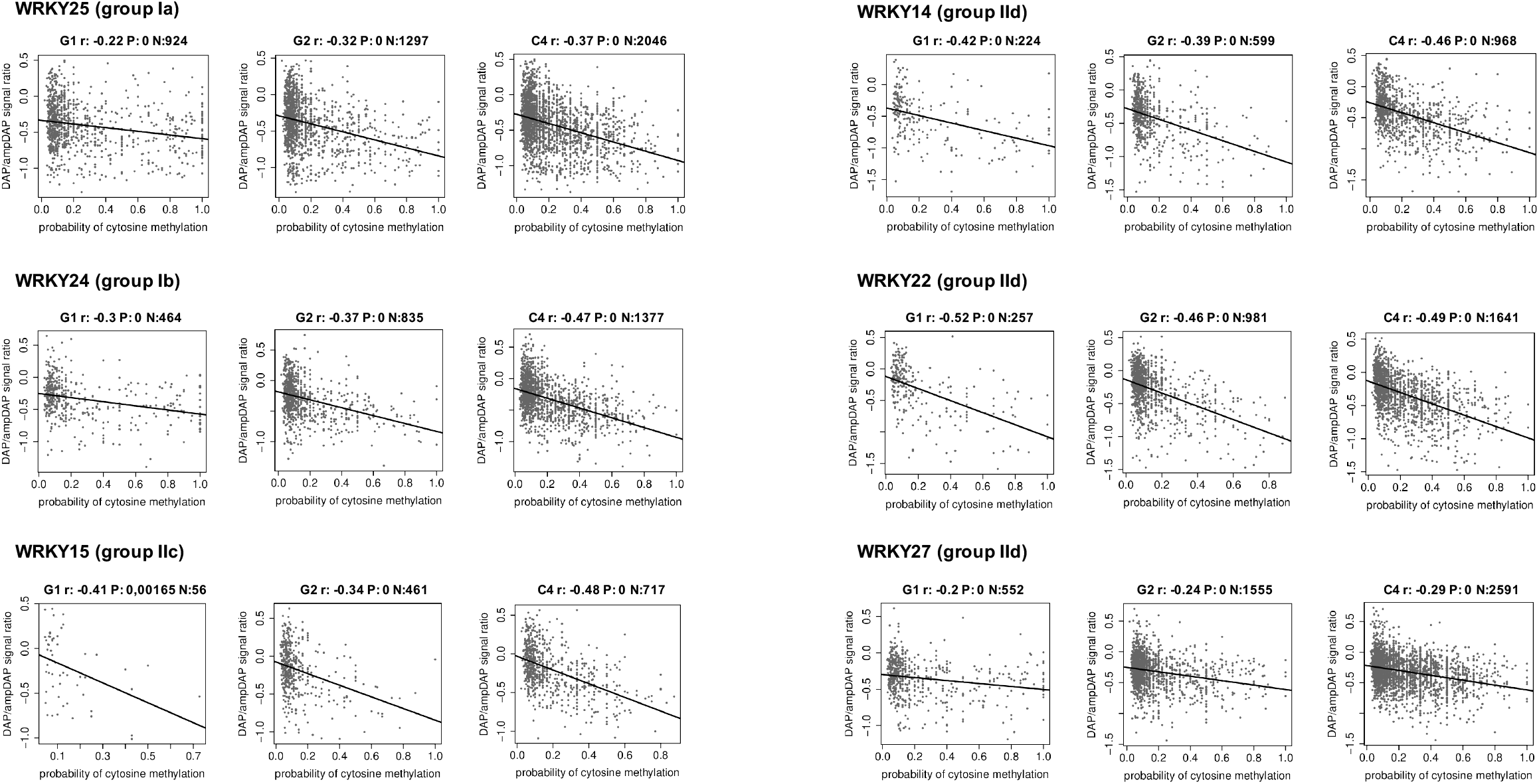
Effect of methylation at each cytosine in the core W- box elements on the DNA binding of AtWRKYs. Effect of methylation on individual positions at the core W-box on the binding of each indicated AtWRKY. Relation between methylation probability at a single nucleotide position in the best predicted WRKY-binding site within bound regions, and the log10-scaled relative binding intensity of a DAP-seq *versus* an ampDAP-seq experiment at bound regions for indicated WRKYs at the 3 different cytosine sites. P-values are adjusted for multiple testing using the Benjamini and Hochberg procedure.

Figure 2 – source data 1. Methylation at the cytosine C4 embedded in a functional W-box has a strong negative impact on AtWRKY40-DNA binding.

**Figure 2 - Supplementary Figure 1.**
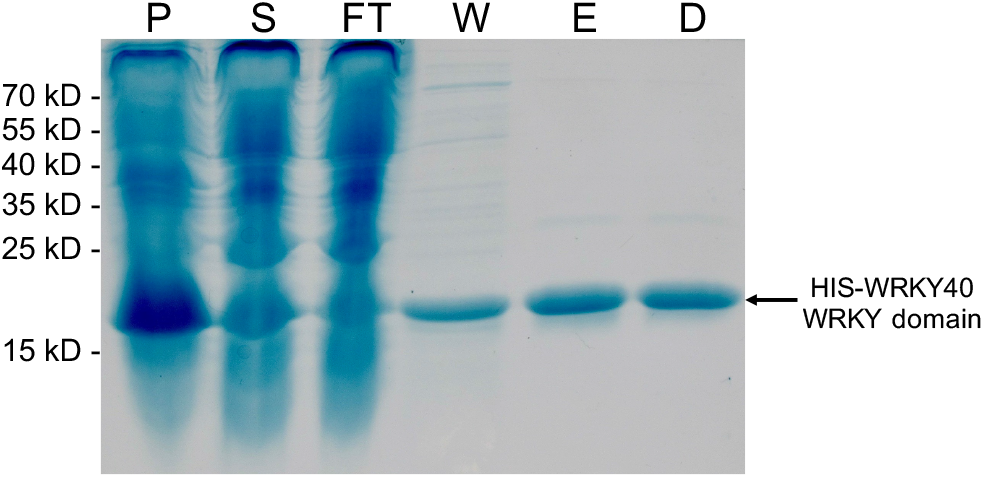
Purification of WRKY40 WRKY domain. Coomassie- blue stained gel showing protein content at different steps of WRKY40 WRKY domain purification. P: Pellet after sonication; S: Supernatant; FT: Supernatant after incubation with Ni- NTA beads; W: Supernatant after washing; E: Elution; D: Elution after dialysis. An arrow indicates the His-WRKY40 WRKY domain.

Figure 2 – Supplementary Figure 1 – source data 1

Coomassie blue stained gel showing purification of WRKY40 WRKY domain

Figure 2 – Supplementary Figure 1 – source data 2

Original picture of the coomassie blue stained gel of WRKY40 WRKY domain

**Figure 2 - Supplementary Figure 2.**
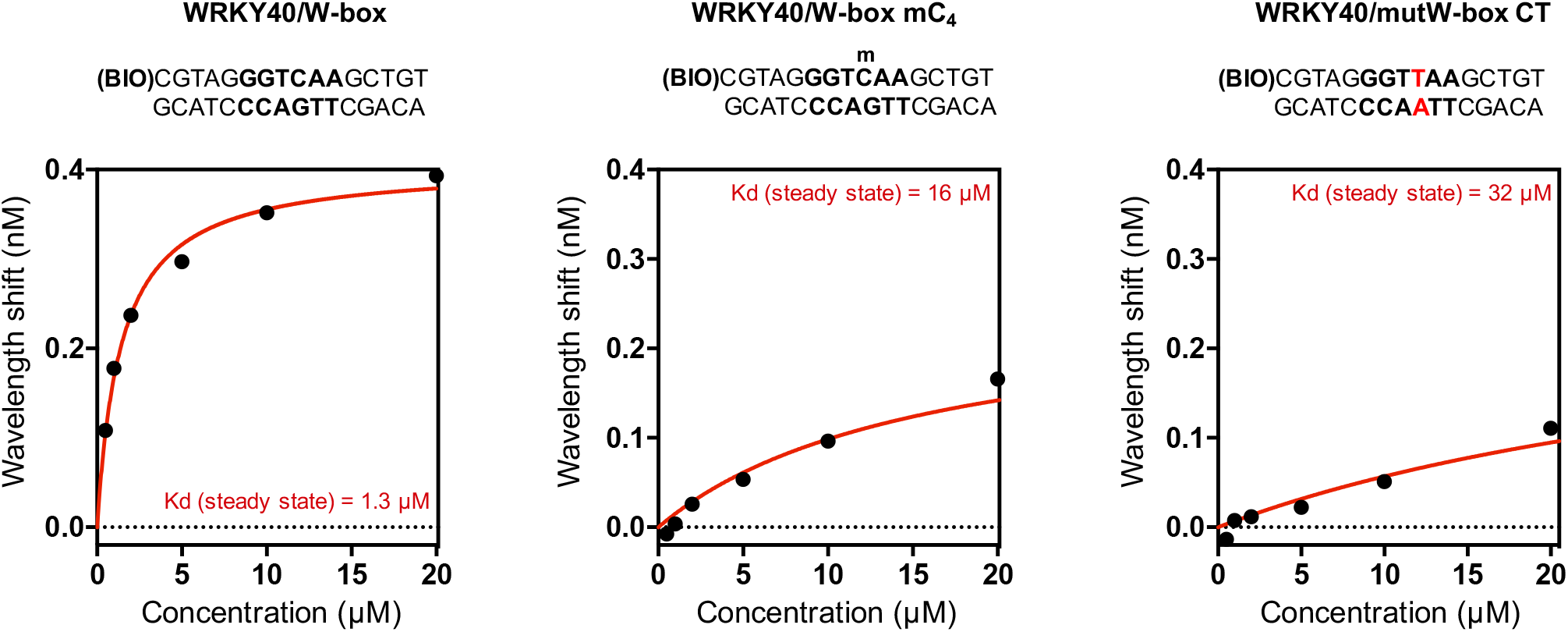
The DNA binding inhibitory effect detected in the presence of 5mC_4_ is almost as strong as the one observed with a point mutation at this specific cytosine. BLI-derived steady state analysis representing binding responses of At- WRKY40 DBD (nM) to DNA duplexes as a function of AtWRKY40 DBD concentration.

Figure 2 – Supplementary Figure 2 – source data 1. The DNA binding inhibitory effect detected in the presence of 5mC_4_ is almost as strong as the one observed with a point mutation at this specific cytosine.

**Figure 3 - Supplementary Figure 1:**
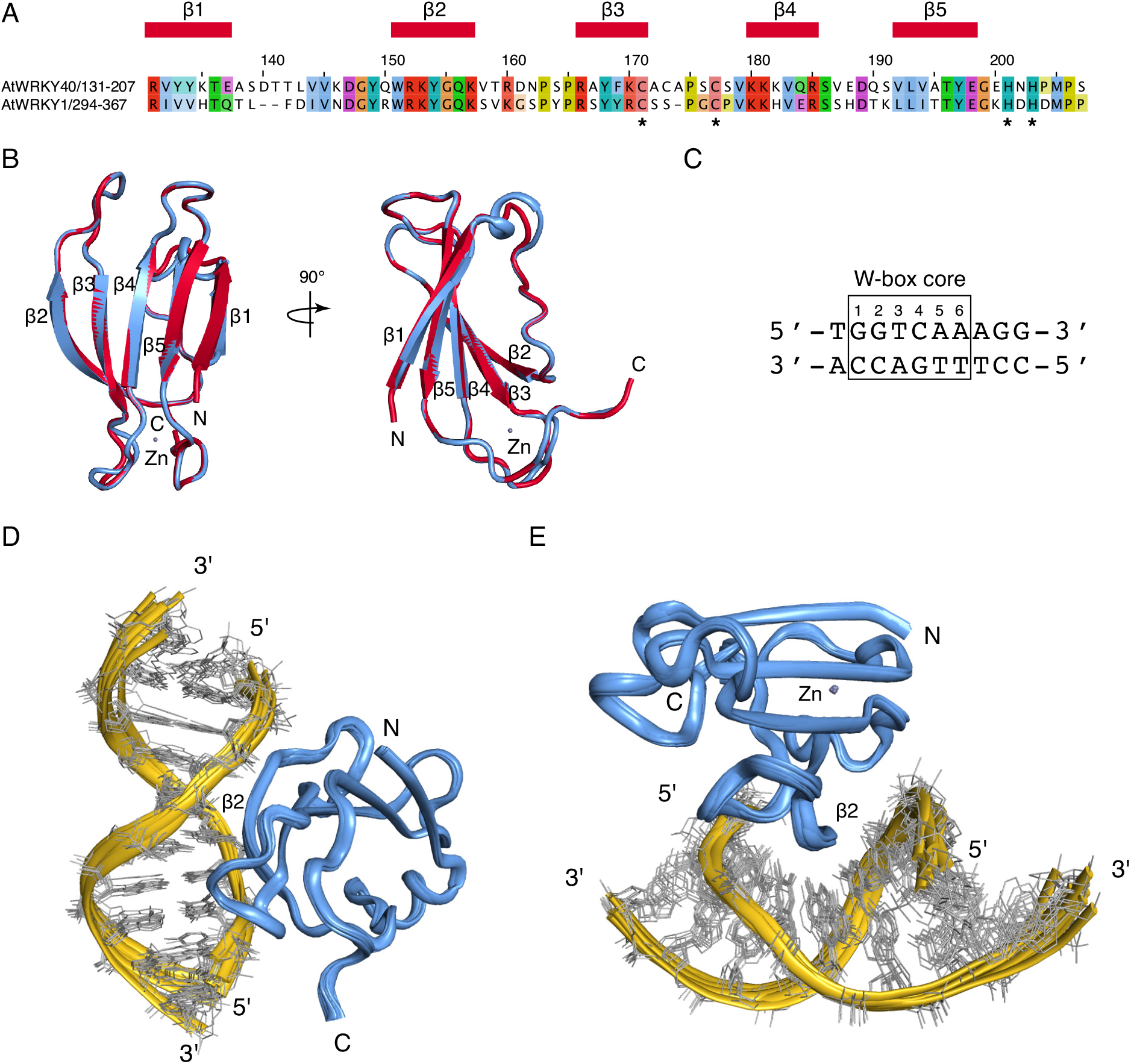
Modelling the interaction of AtWRKY40 with a W- box element. (A) Sequence alignment of the WRKY domain of AtWRKY40 with the C-terminal WRKY domain of AtWRKY1. The alignment is coloured by amino acid conservation and properties. The two domains share 47% of sequence identity. Residue numbering corresponds to that of AtWRKY40. The secondary structure elements of AtWRKY1 (PDB code 2ayd) (Duan et al., 2007) are shown above the alignment. The CCHH zinc-binding motif is indicated by stars below the alignment. (B) Superposition of the C-terminal WRKY domain of AtWRKY1 (in red) with the homology model of the WRKY domain of AtWRKY40 (in light blue). Protein domains are show as cartoons and the five β-strands are labelled. Zinc ions are shown as grey dots. (C) Sequence of the DNA W-box element used for modelling the interaction of AtWRKY40 with DNA. This element corresponds to the W-box element in the structure of the C-terminal WRKY domain of AtWRKY4 in complex with DNA (PDB code 2lex) (Yamasaki et al., 2012). (D, E) Overlay of the 10 lowest energy structures of the AtWRKY40/W- box element model shown in two orientations. The protein is shown as a ribbon in light blue. The DNA is shown in yellow and grey.

**Figure 3 - Supplementary Figure 2:**
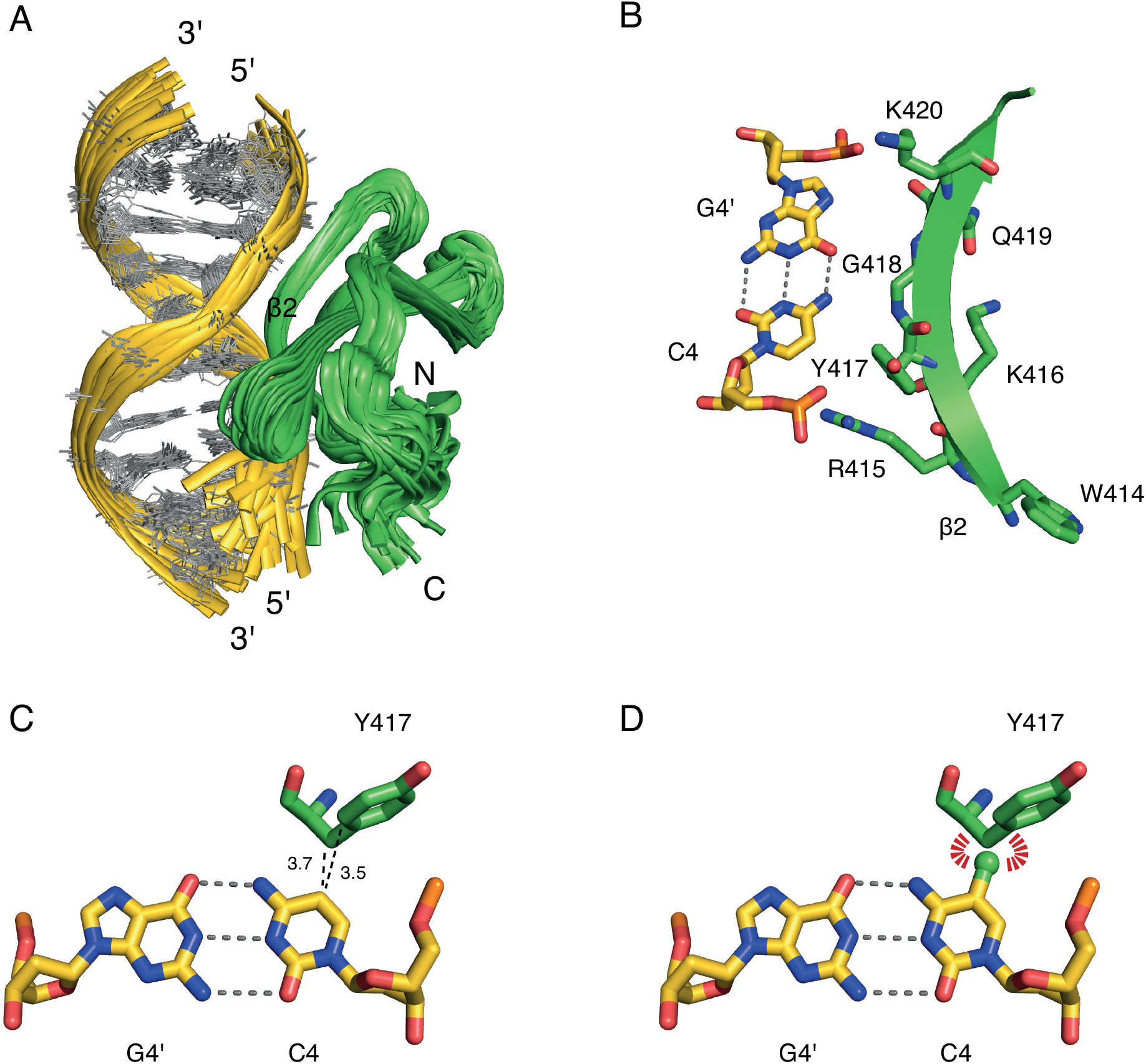
Analysis of the interaction of AtWRKY4 with a W- box element. (A) NMR ensemble of the AtWRKY4/W-box element structure (PDB code 2lex) (Yamasaki et al., 2012). The protein is shown as a ribbon in green. The DNA is shown in yellow and grey. (B) Close-up view of the WRKY motif of β2 (W414-K420). The protein is shown in green with the side chains represented as sticks. The C4-G4’ DNA base-pair is shown as sticks in yellow. The β2 strand enters deeply into the DNA major groove at the level of the C4-G4’ base-pair. In particular, the aromatic ring of C4 makes van der Waals contacts with the side-chain of Y417. (C) The position 5 of C4 makes van der Waals contacts with Y417. (D) Modelling a methyl group (in green) onto unmodified C4 from the W-box core reveals steric hindrance (indicated with red strips) with Y417.

**Figure 3 - Supplementary Figure 3:**
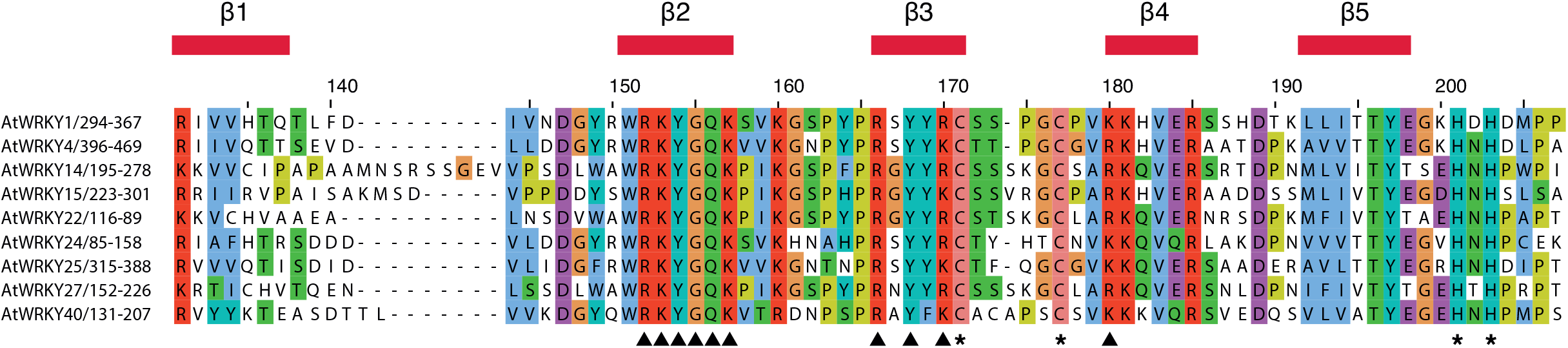
Residues involved in DNA contacts are conserved among different WRKY domains. Sequence alignment of different WRKY domains, namely WRKY domains of AtWRKY1, AtWRKY4, AtWRKY14, AtWRKY15, AtWRKY22, At- WRKY24, AtWRKY25, AtWRKY27 and AtWRKY40. The alignment is coloured by amino acid conservation and properties. Residue numbering corresponds to that of AtWRKY40. The secondary structure elements of AtWRKY1 (PDB code 2ayd) (Duan et al., 2007) are shown above the alignment. The CCHH zinc-binding motif is indicated by stars below the alignment. Residues involved in DNA contacts in both the NMR structure of AtWRKY4 in complex with DNA (PDB code 2lex) and our structural model of AtWRKY40 in complex with DNA (Figure 3) are indicated by arrows below the alignment.

Supplementary file 1. DNA oligonucleotides used in this study

## References

Ando, M., Saito, Y., Xu, G., Bui, N.Q., Medetgul-Ernar, K., Pu, M., Fisch, K., Ren, S., Sakai, A., Fukusumi, T., Liu, C., Haft, S., Pang, J., Mark, A., Gaykalova, D.A., Guo, T., Favorov, A.V., Yegnasubramanian, S., Fertig, E.J., Ha, P., Tamayo, P., Yamasoba, T., Ideker, T., Messer, K., Califano, J.A., 2019. Chromatin dysregulation and DNA methylation at transcription start sites associated with transcriptional repression in cancers. Nat Commun 10, 2188.

Barraud, P., Banerjee, S., Mohamed, W.I., Jantsch, M.F., Allain, F.H.-T., 2014. A bimodular nuclear localization signal assembled via an extended double-stranded RNA-binding domain acts as an RNA-sensing signal for transportin 1. Proceedings of the National Academy of Sciences 111, E1852–E1861.

Barraud, P., Heale, B.S.E., O’Connell, M.A., Allain, F.H.-T., 2012. Solution structure of the N-terminal dsRBD of Drosophila ADAR and interaction studies with RNA. Biochimie 94, 1499–1509.

Birkenbihl, R.P., Kracher, B., Roccaro, M., Somssich, I.E., 2017. Induced Genome-Wide Binding of Three Arabidopsis WRKY Transcription Factors during Early MAMP-Triggered Immunity. Plant Cell 29, 20–38.

Birkenbihl, R.P., Kracher, B., Ross, A., Kramer, K., Finkemeier, I., Somssich, I.E., 2018. Principles and characteristics of the Arabidopsis WRKY regulatory network during early MAMP-triggered immunity. The Plant Journal 96, 487–502.

Brunger, A.T., 2007. Version 1.2 of the Crystallography and NMR system. Nat Protoc 2, 2728–2733.

Campanero, M.R., Armstrong, M.I., Flemington, E.K., 2000. CpG methylation as a mechanism for the regulation of E2F activity. Proceedings of the National Academy of Sciences 97, 6481–6486.

Chen, C., Chen, Z., 2000. Isolation and characterization of two pathogen-and salicylic acid-induced genes encoding WRKY DNA-binding proteins from tobacco. Plant Mol Biol 42, 387–396.

Ciolkowski, I., Wanke, D., Birkenbihl, R.P., Somssich, I.E., 2008. Studies on DNA-binding selectivity of WRKY transcription factors lend structural clues into WRKY-domain function. Plant Mol Biol 68, 81–92.

Clapier, C.R., Cairns, B.R., 2009. The Biology of Chromatin Remodeling Complexes. Annual Review of Biochemistry 78, 273–304.

Comb, M., Goodman, H.M., 1990. CpG methylation inhibits proenkephalin gene expression and binding of the transcription factor AP-2. Nucleic Acids Res 18, 3975–3982.

Cormack, R.S., Eulgem, T., Rushton, P.J., Köchner, P., Hahlbrock, K., Somssich, I.E., 2002. Leucine zipper-containing WRKY proteins widen the spectrum of immediate early elicitor-induced WRKY transcription factors in parsley. Biochimica et Biophysica Acta (BBA)-Gene Structure and Expression 1576, 92–100.

de Pater, S., Greco, V., Pham, K., Memelink, J., Kijne, J., 1996. Characterization of a zinc-dependent transcriptional activator from Arabidopsis. Nucleic Acids Res 24, 4624–4631.

Deleris, A., Halter, T., Navarro, L., 2016. DNA Methylation and Demethylation in Plant Immunity. Annu Rev Phytopathol 54, 579–603.

Dowen, R.H., Pelizzola, M., Schmitz, R.J., Lister, R., Dowen, J.M., Nery, J.R., Dixon, J.E., Ecker, J.R., 2012. Widespread dynamic DNA methylation in response to biotic stress. Proc Natl Acad Sci U S A 109, E2183–2191.

Duan, M.-R., Nan, J., Liang, Y.-H., Mao, P., Lu, L., Li, L., Wei, C., Lai, L., Li, Y., Su, X.-D., 2007. DNA binding mechanism revealed by high resolution crystal structure of Arabidopsis thaliana WRKY1 protein. Nucleic Acids Res 35, 1145–1154.

Eulgem, T., Rushton, P.J., Robatzek, S., Somssich, I.E., 2000. The WRKY superfamily of plant transcription factors. Trends Plant Sci 5, 199–206.

Eulgem, T., Somssich, I.E., 2007. Networks of WRKY transcription factors in defense signaling. Curr Opin Plant Biol 10, 366–371.

Gaston, K., Fried, M., 1995. CpG methylation has differential effects on the binding of YY1 and ETS proteins to the bi-directional promoter of the Surf-1 and Surf-2 genes. Nucleic Acids Res 23, 901–909.

Guertin, M.J., Lis, J.T., 2010. Chromatin Landscape Dictates HSF Binding to Target DNA Elements. PLOS Genetics 6, e1001114.

Halter, T., Wang, J., Amesefe, D., Lastrucci, E., Charvin, M., Singla Rastogi, M., Navarro, L., 2021. The Arabidopsis active demethylase ROS1 cis-regulates defence genes by erasing DNA methylation at promoter-regulatory regions. eLife 10, e62994.

Hinderhofer, K., Zentgraf, U., 2001. Identification of a transcription factor specifically expressed at the onset of leaf senescence. Planta 213, 469–473.

Huang, M., Zhang, Y., Wang, Y., Xie, J., Cheng, J., Fu, Y., Jiang, D., Yu, X., Li, B., 2022. Active DNA demethylation regulates MAMP-triggered immune priming in Arabidopsis. Journal of Genetics and Genomics.

Iguchi-Ariga, S.M., Schaffner, W., 1989. CpG methylation of the cAMP-responsive enhancer/promoter sequence TGACGTCA abolishes specific factor binding as well as transcriptional activation. Genes Dev 3, 612–619.

Iwafuchi-Doi, M., Zaret, K.S., 2016. Cell fate control by pioneer transcription factors. Development 143, 1833–1837.

Iwafuchi-Doi, M., Zaret, K.S., 2014. Pioneer transcription factors in cell reprogramming. Genes Dev. 28, 2679–2692.

Jalili, V., Matteucci, M., Masseroli, M., Morelli, M.J., 2018. Using combined evidence from replicates to evaluate ChIP-seq peaks. Bioinformatics 34, 2338.

Jin, R., Klasfeld, S., Zhu, Y., Fernandez Garcia, M., Xiao, J., Han, S.-K., Konkol, A., Wagner, D., 2021. LEAFY is a pioneer transcription factor and licenses cell reprogramming to floral fate. Nat Commun 12, 626.

Johnson, C.S., Kolevski, B., Smyth, D.R., 2002. TRANSPARENT TESTA GLABRA2, a trichome and seed coat development gene of Arabidopsis, encodes a WRKY transcription factor. Plant Cell 14, 1359–1375.

Khouider, S., Borges, F., LeBlanc, C., Ungru, A., Schnittger, A., Martienssen, R., Colot, V., Bouyer, D., 2021. Male fertility in Arabidopsis requires active DNA demethylation of genes that control pollen tube function. Nat Commun 12, 410.

Kim, J.-S., Lim, J.Y., Shin, H., Kim, B.-G., Yoo, S.-D., Kim, W.T., Huh, J.H., 2019. ROS1-Dependent DNA Demethylation Is Required for ABA-Inducible NIC3 Expression. Plant Physiol 179, 1810–1821.

Klemm, S L., Shipony, Z., Greenleaf, W J., 2019. Chromatin accessibility and the regulatory epigenome. Nat Rev Genet 20, 207–220.

Lai, X., Blanc-Mathieu, R., GrandVuillemin, L., Huang, Y., Stigliani, A., Lucas, J., Thévenon, E., Loue-Manifel, J., Turchi, L., Daher, H., Brun-Hernandez, E., Vachon, G., Latrasse, D., Benhamed, M., Dumas, R., Zubieta, C., Parcy, F., 2021. The LEAFY floral regulator displays pioneer transcription factor properties. Mol Plant 14, 829–837.

Langmead, B., Salzberg, S.L., 2012. Fast gapped-read alignment with Bowtie 2. Nat Methods 9, 357–359.

Law, J.A., Jacobsen, S.E., 2010. Establishing, maintaining and modifying DNA methylation patterns in plants and animals. Nat Rev Genet 11, 204–220.

Le, T.-N., Schumann, U., Smith, N.A., Tiwari, S., Au, P.C.K., Zhu, Q.-H., Taylor, J.M., Kazan, K., Llewellyn, D.J., Zhang, R., Dennis, E.S., Wang, M.-B., 2014. DNA demethylases target promoter transposable elements to positively regulate stress responsive genes in Arabidopsis. Genome Biol. 15, 458.

Lee, M.M., Schiefelbein, J., 1999. WEREWOLF, a MYB-related protein in Arabidopsis, is a position-dependent regulator of epidermal cell patterning. Cell 99, 473–483.

López Sánchez, A., Stassen, J.H.M., Furci, L., Smith, L.M., Ton, J., 2016. The role of DNA (de)methylation in immune responsiveness of Arabidopsis. Plant J. 88, 361–374.

Luo, M., Dennis, E.S., Berger, F., Peacock, W.J., Chaudhury, A., 2005. MINISEED3 (MINI3), a WRKY family gene, and HAIKU2 (IKU2), a leucine-rich repeat (LRR) KINASE gene, are regulators of seed size in Arabidopsis. Proceedings of the National Academy of Sciences 102, 17531–17536.

Machanick, P., Bailey, T.L., 2011. MEME-ChIP: motif analysis of large DNA datasets. Bioinformatics 27, 1696–1697.

Maeo, K., Hayashi, S., Kojima-Suzuki, H., Morikami, A., Nakamura, K., 2001. Role of conserved residues of the WRKY domain in the DNA-binding of tobacco WRKY family proteins. Biosci Biotechnol Biochem 65, 2428–2436.

Magnani, L., Ballantyne, E.B., Zhang, X., Lupien, M., 2011. PBX1 Genomic Pioneer Function Drives ERα Signaling Underlying Progression in Breast Cancer. PLOS Genetics 7, e1002368.

O’Malley, R.C., Huang, S.-S.C., Song, L., Lewsey, M.G., Bartlett, A., Nery, J.R., Galli, M., Gallavotti, A., Ecker, J.R., 2016. Cistrome and Epicistrome Features Shape the Regulatory DNA Landscape. Cell 165, 1280–1292.

Pacis, A., Mailhot-Léonard, F., Tailleux, L., Randolph, H.E., Yotova, V., Dumaine, A., Grenier, J.-C., Barreiro, L.B., 2019. Gene activation precedes DNA demethylation in response to infection in human dendritic cells. Proc. Natl. Acad. Sci. U.S.A. 116, 6938–6943.

Pacis, A., Tailleux, L., Morin, A.M., Lambourne, J., MacIsaac, J.L., Yotova, V., Dumaine, A., Danckaert, A., Luca, F., Grenier, J.-C., Hansen, K.D., Gicquel, B., Yu, M., Pai, A., He, C., Tung, J., Pastinen, T., Kobor, M.S., Pique-Regi, R., Gilad, Y., Barreiro, L.B., 2015. Bacterial infection remodels the DNA methylation landscape of human dendritic cells. Genome Res. 25, 1801–1811.

Qiu, Q., Mei, H., Deng, X., He, K., Wu, B., Yao, Q., Zhang, J., Lu, F., Ma, J., Cao, X., 2019. DNA methylation repels targeting of Arabidopsis REF6. Nat Commun 10, 2063.

Rushton, P.J., Torres, J.T., Parniske, M., Wernert, P., Hahlbrock, K., Somssich, I.E., 1996. Interaction of elicitor-induced DNA-binding proteins with elicitor response elements in the promoters of parsley PR1 genes. EMBO J 15, 5690–5700.

Schumann, U., Lee, J.M., Smith, N.A., Zhong, C., Zhu, J.-K., Dennis, E.S., Millar, A.A., Wang, M.-B., 2019. DEMETER plays a role in DNA demethylation and disease response in somatic tissues of Arabidopsis. Epigenetics 14, 1074–1087.

Tate, P.H., Bird, A.P., 1993. Effects of DNA methylation on DNA-binding proteins and gene expression. Curr Opin Genet Dev 3, 226–231.

Ulker, B., Somssich, I.E., 2004. WRKY transcription factors: from DNA binding towards biological function. Curr Opin Plant Biol 7, 491–498.

Wang, B., Luo, Q., Li, Y., Yin, L., Zhou, N., Li, X., Gan, J., Dong, A., 2020. Structural insights into target DNA recognition by R2R3-MYB transcription factors. Nucleic Acids Research 48, 460–471.

Wang, Z., Yang, P., Fan, B., Chen, Z., 1998. An oligo selection procedure for identification of sequence-specific DNA-binding activities associated with the plant defence response. Plant J 16, 515–522.

Waterhouse, A., Bertoni, M., Bienert, S., Studer, G., Tauriello, G., Gumienny, R., Heer, F.T., de Beer, T.A.P., Rempfer, C., Bordoli, L., Lepore, R., Schwede, T., 2018. SWISS-MODEL: homology modelling of protein structures and complexes. Nucleic Acids Res 46, W296–W303.

Watt, F., Molloy, P.L., 1988. Cytosine methylation prevents binding to DNA of a HeLa cell transcription factor required for optimal expression of the adenovirus major late promoter. Genes Dev 2, 1136–1143.

Yamaguchi, N., 2021. LEAFY, a Pioneer Transcription Factor in Plants: A Mini-Review. Frontiers in Plant Science 12.

Yamamuro, C., Miki, D., Zheng, Z., Ma, J., Wang, J., Yang, Z., Dong, J., Zhu, J.-K., 2014. Overproduction of stomatal lineage cells in Arabidopsis mutants defective in active DNA demethylation. Nat Commun 5, 4062.

Yamasaki, K., Kigawa, T., Watanabe, S., Inoue, M., Yamasaki, T., Seki, M., Shinozaki, K., Yokoyama, S., 2012. Structural basis for sequence-specific DNA recognition by an Arabidopsis WRKY transcription factor. J Biol Chem 287, 7683–7691.

Yin, Y., Morgunova, E., Jolma, A., Kaasinen, E., Sahu, B., Khund-Sayeed, S., Das, P.K., Kivioja, T., Dave, K., Zhong, F., Nitta, K.R., Taipale, M., Popov, A., Ginno, P.A., Domcke, S., Yan, J., Schübeler, D., Vinson, C., Taipale, J., 2017. Impact of cytosine methylation on DNA binding specificities of human transcription factors. Science 356, eaaj2239.

Yu, A., Lepère, G., Jay, F., Wang, J., Bapaume, L., Wang, Y., Abraham, A.-L., Penterman, J., Fischer, R.L., Voinnet, O., Navarro, L., 2013. Dynamics and biological relevance of DNA demethylation in Arabidopsis antibacterial defense. Proceedings of the National Academy of Sciences 110, 2389–2394.

Zaret, K.S., 2020. Pioneer Transcription Factors Initiating Gene Network Changes. Annual Review of Genetics 54, 367–385.

Zhang, Q., Wang, D., Lang, Z., He, L., Yang, L., Zeng, L., Li, Y., Zhao, C., Huang, H., Zhang, Heng, Zhang, Huiming, Zhu, J.-K., 2016. Methylation interactions in Arabidopsis hybrids require RNA-directed DNA methylation and are influenced by genetic variation. Proc Natl Acad Sci U S A 113, E4248–4256.

Zhang, X., Yazaki, J., Sundaresan, A., Cokus, S., Chan, S.W.-L., Chen, H., Henderson, I.R., Shinn, P., Pellegrini, M., Jacobsen, S.E., Ecker, J.R., 2006. Genome-wide High-Resolution Mapping and Functional Analysis of DNA Methylation in Arabidopsis. Cell 126, 1189–1201.

